# Bayesian inference in arm posture perception

**DOI:** 10.1101/2024.07.05.602180

**Authors:** Valeria C. Peviani, Manon G.A. Joosten, Luke E. Miller, W. Pieter Medendorp

## Abstract

To configure our limbs in space the brain must compute their position based on sensory information provided by mechanoreceptors in the skin, muscles, and joints. Because this information is corrupted by noise, the brain is thought to process it probabilistically, and integrate it with prior belief about arm posture, following Bayes’ rule. Here, we combined computational modeling with behavioral experimentation to test this hypothesis. The model conceives the perception of arm posture as the combination of a probabilistic kinematic chain composed by the shoulder, elbow, and wrist angles, compromised with additive Gaussian noise, with a Gaussian prior about these joint angles. We tested whether the model explains errors in a VR-based posture-matching task better than a model that assumes a uniform prior. Human participants (N=20) were required to align their unseen right arm to a target posture, presented as a visual configuration of the arm in the horizontal plane. Results show idiosyncratic biases in how participants matched their unseen arm to the target posture. We used maximum likelihood estimation to fit the Bayesian model to these observations and retrieve key parameters including the prior means and its variance-covariance structure. The Bayesian model including a Gaussian prior explained the response biases and variance much better than a model with a uniform prior. The prior varied across participants, consistent with the idiosyncrasies in arm posture perception, and in alignment with previous behavioral research. Our work clarifies the biases in arm posture perception within a new perspective on the nature of proprioceptive computations.

**New & Noteworthy:** We modeled the perception of arm posture as a Bayesian computation. A VR posture-matching task was used to empirically test this Bayesian model. The Bayesian model including a non-uniform postural prior well explained individual participants’ biases in arm posture perception.

## Introduction

Proprioception is defined as the awareness of the mechanical and spatial state of the body and its musculoskeletal parts (1). It is essential in several of our functions and behaviors, such as the planning and execution of our movements, the perception of force and weight, as well as the notion of our body image and the perception of self (2). As a sense, proprioception is mediated not only by mechanoreceptors in the skin, muscle, tendons, and joints, but also by predictions derived from efferent motor commands. For instance, to determine the pose of the arm in three-dimensional space, the brain needs to combine feedback and forward signals informing about flexion of the shoulder, elbow, and wrist (3, 4).

Different tasks are used to assess the acuity of arm proprioception, including the posture matching task (5). In this task, a participant is asked to match the unseen arm to a set of target joint angles. The resulting proprioceptive estimates are characterized by both systematic and variable errors that vary across arm postures (6–8); for instance, the elbow angle is perceived as more flexed for extended postures (9–11).

Several explanations have been hypothesized for these errors, ranging from central adaptation in the processing of peripheral signals (5), asymmetries in the distribution of preferred directions of muscle spindles (12), to a decay of proprioceptive memory (6). It has also been proposed that proprioceptive estimates are biased towards ‘comfortable’ (13) or ‘default’ (14–16) postures, which may be based on the most frequently adopted arm positions (17, 18). Apart from these more heuristic explanations, there is no model that formalizes the neural computations underlying arm posture perception and can provide a unifying explanation for the observed systematic and variable errors. This is the goal of the present study.

To develop this model, we view the arm as a probabilistic kinematic chain of joint angles, whose proprioceptive measurements are contaminated by noise (7, 19–21). As formalized within the Bayesian framework (22, 23), the brain may simulate this probabilistic chain by forming a belief (the posterior) about the most probable arm pose by integrating the noisy proprioceptive measurements (the likelihood) with prior beliefs, which are possibly inferred from an accumulated history of previous arm postures. The final posterior is the weighted average of the likelihood and prior, with the weights proportional to the inverse of their variance. The effect of the prior amounts to an increase in systematic errors (i.e., a bias) but a decrease in the variable errors in arm posture perception.

To test this model empirically, we designed a VR-based proprioceptive matching task in which participants were required to align their unseen arm to various target postures, presented as a visual configuration of the arm in the horizontal 2D plane. For each participant, we found systematic and variable errors in the perceived orientation of the shoulder, elbow and wrist joints that could be well explained by the Bayesian integration of proprioceptive information and a postural prior. Our work clarifies the biases in arm posture perception within a new perspective on the computations in proprioceptive processing.

## Materials and Methods

### Computational model

Configuring the arm in a certain posture requires adjusting the angles of three joints (shoulder, elbow, and wrist) connecting the upper arm, the forearm, and the hand. If we consider the horizontal plane, the physical orientation of the arm can thus be defined as a set of 1D joint angles **θ** = [*θ*_*s*_, *θ*_*e*_, *θ*_*w*_], corresponding to the shoulder, elbow, and wrist angle, respectively (**Figure 1A**), where **θ** = [0°, 0°, 0°] defines a fully extended arm parallel to the torso.

**Figure 1.**
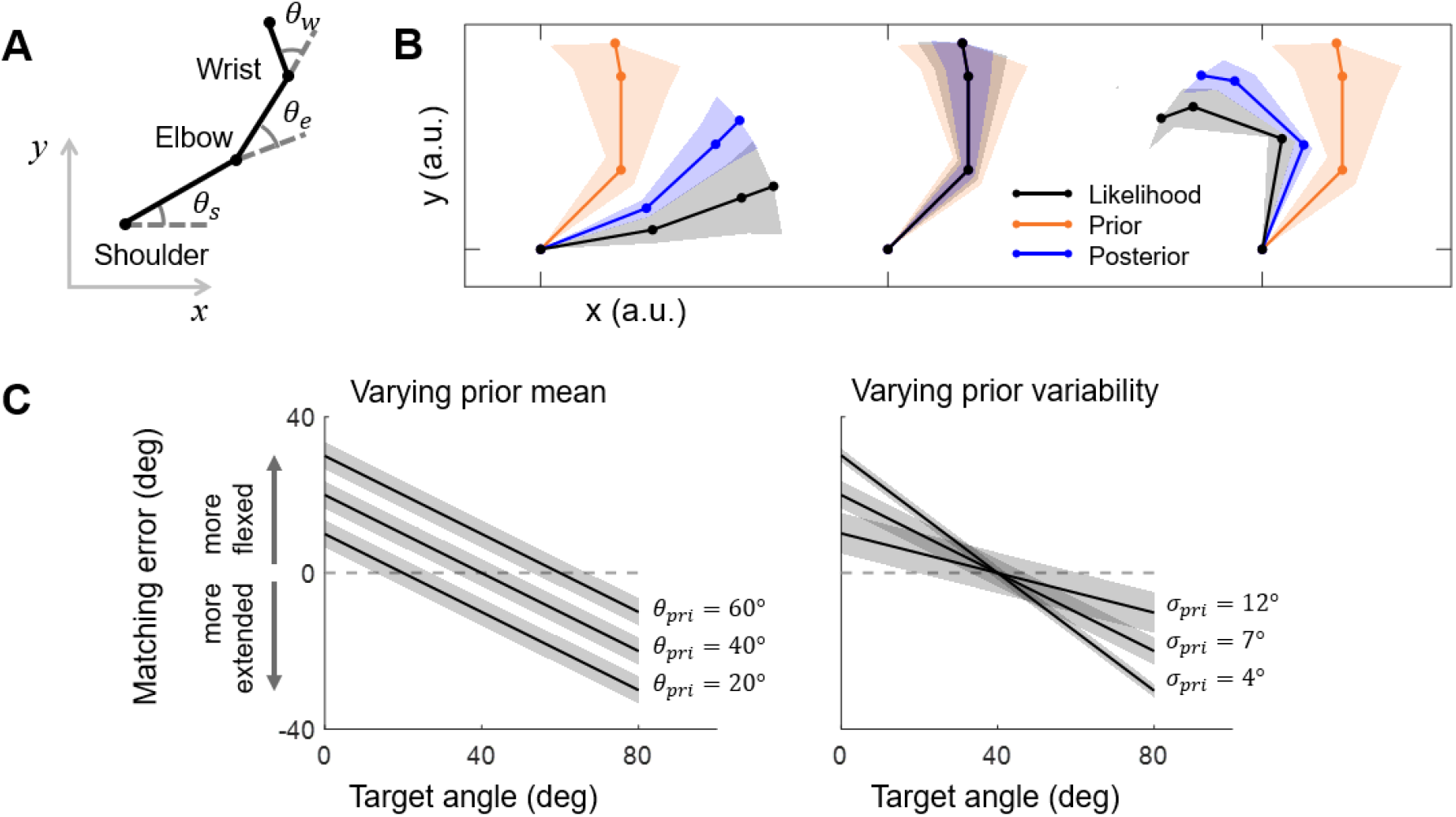
Model predictions. **A**. Kinematic chain representing the arm in a 2D plane as a set of joint angles (shoulder, elbow, and wrist). **B**. Bayesian integration in arm posture perception. The prior is shown in orange. The posterior (in blue) is shown for three different arm postures (in black). Shaded area represents standard deviation. While the model performs the Bayesian integration in angular coordinates, the effects are shown in Cartesian coordinates, including the error propagation due to the kinematic chain. **C**. Matching errors (perceived – real angle) as a function of joint angle for a single joint for three different prior means, i.e., ***μ***_*pr*_ (left panel) and prior variability, i.e., *σ*_*lik*_ (right panel). The dashed line shown the model prediction with a uniform prior.

We assumed that the proprioceptive signals about joint angles are accurately calibrated (i.e., unbiased) but corrupted by independent Gaussian noise. Here, we applied Bayes’ rule to estimate arm posture from these signals. For simplicity, we approximated the likelihood distribution, provided by the proprioceptive inputs, as well as the prior as Gaussian probability distributions. Of note, since the standard deviations of these Gaussians are restricted to moderate levels, we refrained from using their circular analog (Von Mises distribution) as they provided analytically non-tractable solutions.

The likelihood is thus specified as a Gaussian distribution with its mean ***μ***_*lik*_ defined by the involved joint angles (*θ*_*s*_, *θ*_*e*_, *θ*_*w*_); its variability *∑*_*lik*_ is defined as a diagonal variance-covariance matrix specifying variance of the respective joint angles 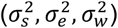 and off-diagonal values set to zero (assuming uncorrelated noise). The prior is also defined as a Gaussian distribution in joint coordinates with mean ***μ***_*pr*_ and diagonal variance-covariance matrix ∑_*pr*_, allowing covariance between subsequent joint angles (i.e., between *θ*_*s*_ and *θ*_*e*_, *θ*_*e*_ and *θ*_*w*_). The posterior (***μ***_*pos*_, ∑_*pos*_) is given by the weighted average of the likelihood and prior, following:

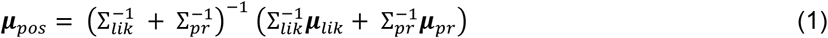

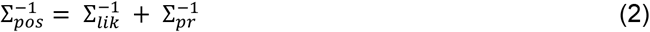

Note that eq. (2) represents ∑_*pos*_ on a single trial, i.e., the variance of the posterior distribution on a given trial. However, in an experiment, we measure the variability of behavioral responses across several trials, i.e., the variance of response distribution (∑_*r*_) (24, 25). The variance of the response distribution can be computed as:

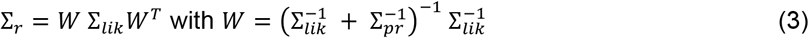

If the likelihood and prior are assumed to be Gaussians with a constant standard deviation, irrespective of arm posture, the response distribution will be Gaussian as well. Under these the key prediction of the model is a bias in posture matching which changes across different arm postures (**Figure 1B, 1C**). The bias is directed towards the prior. As shown, matching errors (response angle – target angle) for each joint will be negative (the joint is perceived as more extended) if the actual arm posture is more flexed than the prior posture and negative (the joint is perceived as more flexed) if the actual posture is more extended than the prior posture. A model with a uniform prior would instead predict no biases (dashed gray line in **Figure 1C**).

To test this model empirically, we designed a proprioceptive matching task (described in detail below) to measure how well participant can their shoulder, elbow and wrist orientation to various target joint angles.

### Participants

Twenty-one (17 females) healthy participants were recruited for the experiment. Their mean age (±SD) was 24.1 ± 2.8 yrs. Participants had normal or corrected-to-normal vision and were all right-handed, as assessed by the Edinburgh Handedness Inventory, Short Form (26, 27), with laterality quotient 91.4 ± 11.8 (mean±SD). The study was approved by Ethics Committee of the Faculty of Social Sciences of the Radboud University. All participants gave their written informed consent. The data collection of one participant was not completed due to discomfort with the task (final N = 20).

### Setup

Participants were seated at a table whose surface was adjusted to the level of their shoulder, ensuring that their right arm only moved in the horizontal plane. Participants had their right forearm and right hand resting on air sleds to minimize movement friction (**Figure 2A**), and their shoulder was fixed in place by a custom-made support.

**Figure 2.**
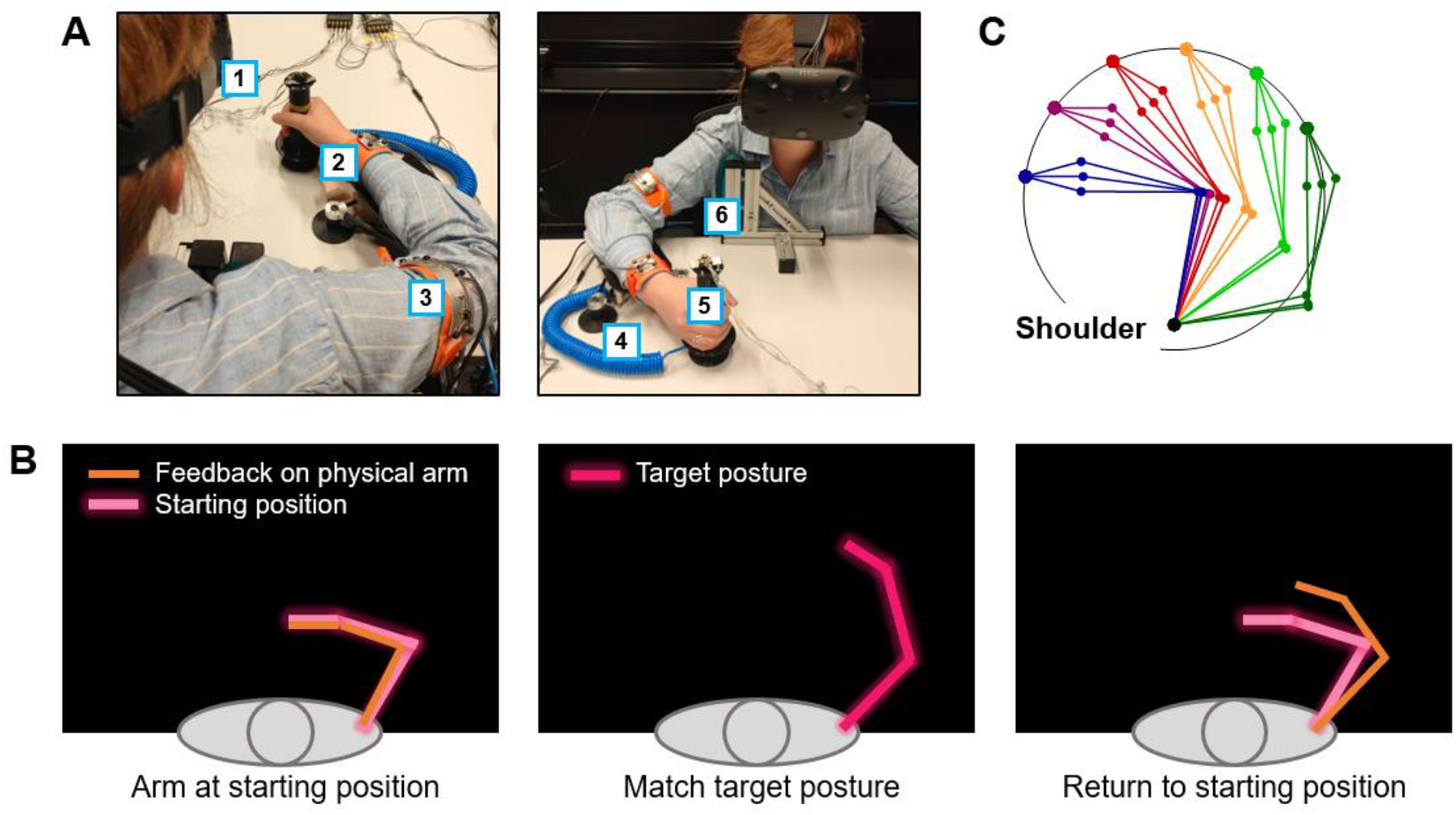
Experiment. **A**. Experimental set-up: participants were seated at a table with the VR headset on (1). Infrared markers configured in custom-made rigid bodies (e.g., 2,3) were used to measure the position and orientation of the right shoulder, elbow, and wrist. Participants rested their forearm and hand on supporting airsleds (4,5) to minimize friction during the arm movement. The position of the shoulder was fixed by a custom-made support (6). **B**. Trial configuration, seen from a top view: At the beginning of each trial, participants brought their physical arm in visual alignment with the starting posture (left panel). Next, participants had to align their unseen arm with the target posture (middle panel) and return their arm to the starting position. **C**. Target postures were calculated based on the individual arm size, so that the endpoint fell on a semi-circle with 30 cm radius.

We used infrared motion tracking (100 Hz sampling rate; Optotrak Certus, Northern Digital Inc.) to track the position of markers placed on the right arm. First Principles software (Northern Digital Inc.) was used to align coordinate systems of all position sensors into a global room-based coordinate system (i.e., registration). We attached custom-made rigid bodies to each individual limb segment (upper arm, forearm, and hand) to determine real-time the position and orientation of the shoulder, elbow, and wrist. Using these measures, we calculated the length of each limb segment, which served as participant-specific input in the design of the stimuli in the VR task (see (25) for a similar approach).

The VR environment was created with Unity (Unity Technologies) and rendered through an HTC Vive VR headset. The external light of the room was dimmed to prevent peripheral cues around the headset. Images were presented with 1080*1200 pixel resolution to each eye at a 90 Hz refresh rate, creating a virtual field of view of 110°. The VR space was aligned with the Optotrak space using a calibration procedure supported by custom-made Python scripts. The alignment error was within 0.5 and 1.1 mm for all testing sessions. Time stamps from the VR system were used to segment the continuous motion tracking signal into trials.

### Paradigm

We designed a novel VR-based arm proprioceptive matching task in which participants were required to align their unseen right arm (upper arm, forearm, and hand) with a visual stimulus representing a target posture in the 2D horizontal plane. In the VR environment, each target posture was represented in the first-person perspective as three connected lines corresponding to upper arm, forearm, and hand, matching the participant’s actual arm length.

The structure of a trial is represented in **Figure 2B**: At the beginning of each trial, participants had realtime visual feedback of their physical arm, rendered in VR as three segments matching with the physical position and lengths of their arm. Participants were instructed to align their physical arm with a starting posture, also represented as three interconnected lines with angles: shoulder 62.5°, elbow 92.5°, wrist 0°, jittered across trials up to 2.5°, 2.5° and 2°. Once they aligned their arm with the starting posture, they confirmed this by pressing a button on a VR controller held in their left hand. Thereafter, a target posture appeared while the visual feedback of the starting posture disappeared. Participants were then required to align their unseen arm with the target posture, confirmed the response with a button press, and move back to the starting position to await the next trial. Visual feedback about the physical arm posture reappeared two seconds after the response was given.

The joint angles of the target postures were based on the position of six endpoints at a semicircle with 30 cm radius (**Figure 2C**). We used inverse kinematics to derive a set of target postures for each participants given these endpoints, participants’ arm segment lengths, and the following constraints: i) shoulder angle within 0° and 110°; ii) elbow angle within 1° and 110°; iii) wrist angle within - 20° and 45°, iv) a minimum spacing of 10° between each target angle. Due to individual variations in arm size, the number of target postures given those constraints varied across participants (16 to 22 unique postures per participant). Each posture was repeated ten times in a randomized order, for a total of 160 to 220 trials per participant (mode: 180 trials). Trials were self-paced and the task lasted approximately 45 minutes. Before starting the actual experiment, participants received verbal instructions and performed a training session, including twelve trials (six postures were randomly sampled and repeated twice in a random order). During the training session participants continuously had real-time visual feedback about their physical arm position.

### Behavioral analysis

#### Response kinematics

Motion tracking data were processed in MATLAB (R2022b). The continuous motion tracking signal was segmented into trials. Matching responses for each trial and joint (shoulder, elbow, wrist) were computed using inverse kinematics. For descriptive statistics, we computed the matching error in degree for each trial and joint as (response angle – target angle). The following section describes how the model described above was fitted to the observed matching responses, separately to each individual participant.

#### Model fitting

The main model contains nine free parameters (**Table 1**) that were fitted to all data simultaneously for each participant. For the likelihood (***μ***_*lik*_), we assumed that three joint angles were noisy but unbiased. The parameter for the mean of each joint estimate was therefore fixed to its experimental value, i.e., the target joint angle of that trial. The variance of these joint estimates (the diagonal values of the variance-covariance matrix) was free to vary within realistic constraints (**Table 1**) given our experimental design (see above) and previous research (28). We assumed that the variance in these parameters is uncorrelated, such that the off-diagonal values of the variance-covariance matrix were set to zero. In total, the fitting for the likelihood had three free parameters.

**Table 1.** Lower and upper bounds for the model parameters.

|  | Likelihood st. dev. (deg) |  |  | Prior mean (deg) |  |  | Prior st. dev. (deg) |  |  |
| --- | --- | --- | --- | --- | --- | --- | --- | --- | --- |
| | $\sigma_s$ | $\sigma_e$ | $\sigma_w$ | $\theta_s$ | $\theta_e$ | $\theta_w$ | $\sigma_s$ | $\sigma_e$ | $\sigma_w$ |
| **LB** | 0.1 | 0.1 | 0.1 | -5 | 0 | -70 | 0.1 | 0.1 | 0.1 |
| **UB** | 20 | 20 | 20 | 120 | 120 | 70 | 30 | 30 | 30 |

The parameter for the mean and variance of the prior for each joint estimate (***μ***_*pr*_) were free to vary within realistic constraints (**Table 1**). For simplicity off-diagonal values were set at zero in variance-covariance matrix representing the prior. In total, the fitting for the prior had six free parameters.

We also tested three variants of this model, which differed in their assumption of the prior. To test that matching responses are not solely based on sensory inputs, we fitted the same model under the assumption of a uniform prior. This comes down to excluding the prior from the model fitting, which essentially means that this model variant (model variant 1: uniform prior) has three free parameters (the three likelihood variances). Second, we tested whether the prior merely reflects some task contextual features, such as such as the statistics of the target angles (model variant 2: mean target posture prior), or the frequently-held the starting posture (model variant 3: starting posture prior). For these two model variants the mean values of the prior were fixed (mean target posture for model variant 2; starting posture for model variant 2), whereas six free parameters (3 likelihood variances and 3 prior variances) were fit to the data. Lower and upper bounds for parameter search space for the alternative models were the same as the main model (**Table 1**).

To estimate best fitting parameters for each participant, we maximized the probability of the matching responses given a range of initial parameters. We applied Maximum Likelihood Estimation (MLE) to maximize the log-likelihood in relation to parameters of the main model or its variants. This log-likelihood function (LL) was defined as a 3D Gaussian function:

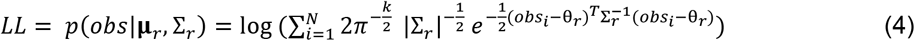

in which N is the number of observations (= n of trials*n of joints); k is the number of dimensions (3); obs represent observed angle responses (in degrees); **μ**_*r*_ and ∑_*r*_ are mean and covariance of the response distribution (in degrees), given each initial set of free parameters. Optimal parameter values were obtained by minimizing the negative likelihood function using the Matlab routine “fmincon”, e.g., (24, 25). Per participant, this model was fitted 100 times. For each fitting run, random values within fixed bounds (**Table 1**) were used as start values for the free parameters to maximize the possibility to find the best fitting values. We then selected the best fit as the one associated with the lowest -LL.

To validate our fitting procedure, we performed a parameter recovery analysis. Using the main generative model, we simulated 100 datasets from sets of known parameters drawn from a continuous uniform distribution having the same bounds as reported in **Table 1**. Next, we fitted the generative model to this data, using 100 parameters initializations for every model fit. **Supplemental Figure 1** shows the scatter plots of the predicted values against the generative values along with the regression line for every model parameter. Parameter recovery was very good with R^2^ > 0.79.

To compare the quality of the main model and its variants, we computed their BIC (Bayesian Information Criterion) scores and averaged across participants. The BIC expresses the maximized log-likelihood (LL) as a function of the number of observations (N) and of free parameters (n):

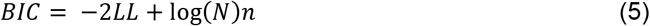

with a lower BIC indicates a better fit. As an indication of the quality of the model fits, we also computed the R^2^ for each participant. This was calculated by as:

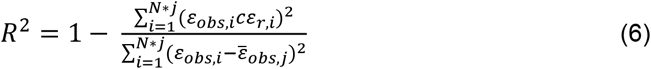

where N is the number of observations, j is the number of joints, and the observed (obs) and predicted errors (*ε*) for each trial and joint are calculated as observed angle – target angle.

## Results

We used a proprioceptive matching task to assess arm posture perception. Using a VR environment, participants were instructed to align their unseen arm with target postures displayed as configurations of visual line segments. We measured how well their shoulder, elbow and wrist angle match the target angles.

**Figure 3A** shows the matching errors (black dots) of the three joints as a function of their target angle for four representative participants. In the same format, **Figure 3B** illustrates the average errors across all 20 participants. For all joints, matching errors varied significantly with target angle (linear regression, all p<.001, not shown), consistent with the predictions of our main model (**Figure 1C**). For the shoulder, average matching errors were overall smaller than for the elbow and wrist (average absolute errors ± SD: 3.4±2.8° versus 5.6±4.6° and 5.8±4.4°) and mostly positive, suggesting that participants perceived it to be more flexed than its actual orientation. For the elbow and wrist, positive and negative errors were observed, suggesting that their perceived position was consistently pulled towards midrange joint angles. For these two joints, matching errors were also more variable than for shoulder (SD is 12.6±3.1° and 11.5±2.3° versus 6.5±1.8°).

**Figure 3.**
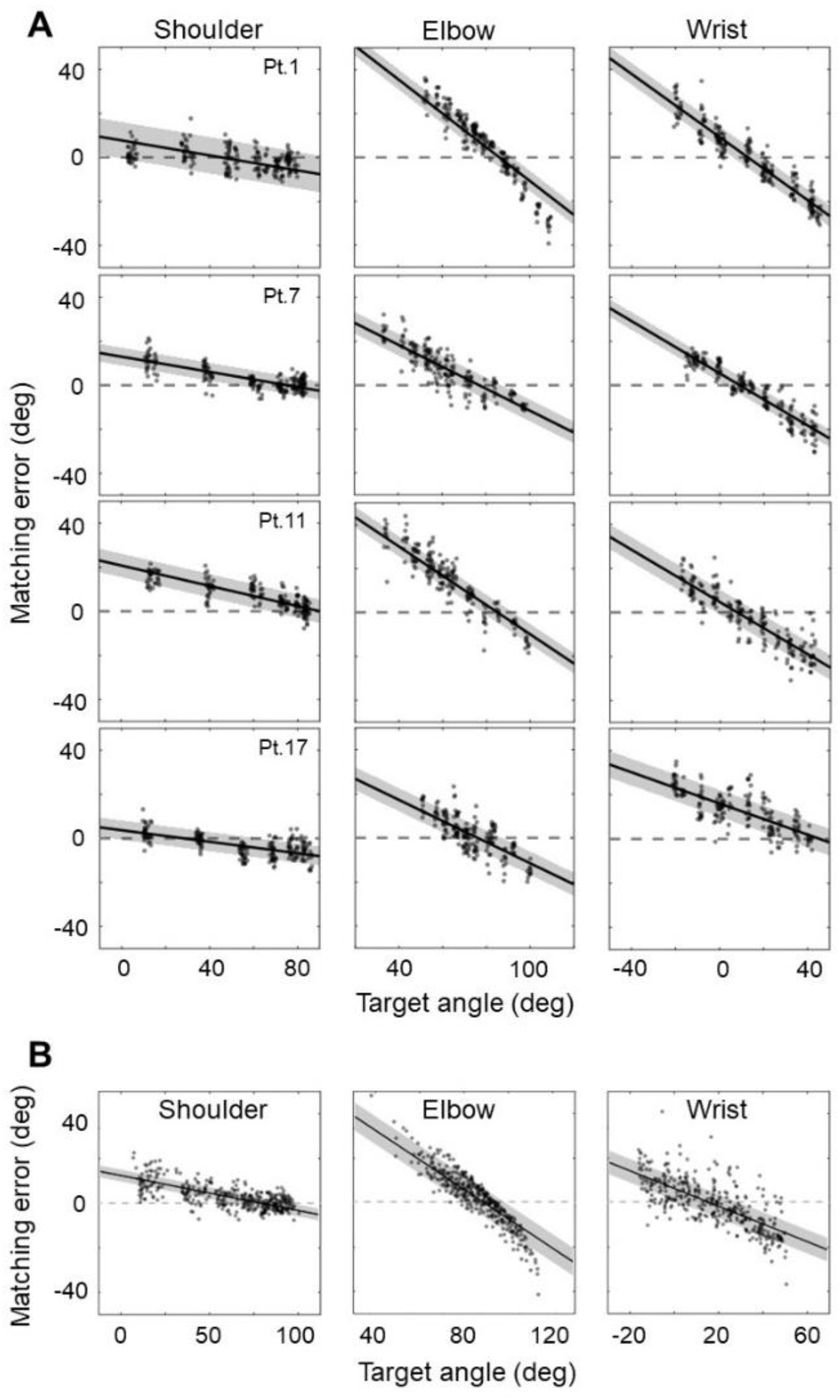
Matching errors as a function of target angle for shoulder, elbow, and wrist. **A**. Four representative participants. **B**. Averaged errors across participants. Lines show predictions of a Bayesian model with a Gaussian postural prior, fit as free parameter. The dashed gray line represents predictions of the model with a uniform prior.

We examined how well our main Bayesian model could account for these observations. **Figure 3** shows the best-fit lines (±SD) of the model for the four representative participants (**Figure 3A**) and as the average of all fit lines (±SD) at the group level (**Figure 3B**). **Table 2** reports best fitted parameter values, for each individual participant along with their group-level average. The quality of the fit, as expressed by R^2^ (last column, **Table 2**), varied between 0.44 and 0.85 (median: 0.72, IQR: 0.16), indicating that the model explains roughly three quarters of the matching response variance.

**Table 2.** Main model fitted parameters and goodness of fit. The best fitting parameters for the likelihood and prior variance, expressed as SD (***σ***_***lik***_, ***σ***_***pr***_) and prior mean (**θ**_***pr***_) pertaining to the shoulder (s), elbow (e) and wrist (w) are reported for each participant. On the rightward part of the table, the minimized negative log-likelihood (eq. 4), BIC (eq. 5) and variance explained by the model as R^2^ (eq. 6).

| Pt. | Best fitting parameters |  |  |  |  |  |  |  |  | Fit outcome |  |  |
| --- | --- | --- | --- | --- | --- | --- | --- | --- | --- | --- | --- | --- |
| | $\sigma_{lik,s}$ | $\sigma_{lik,e}$ | $\sigma_{lik,w}$ | $\theta_{pr,s}$ | $\theta_{pr,e}$ | $\theta_{pr,w}$ | $\sigma_{pr,s}$ | $\sigma_{pr,e}$ | $\sigma_{pr,w}$ | -LL | BIC | R<sup>2</sup> |
| 1 | 10.0 | 20.0 | 17.3 | 45.3 | 86.1 | 12.9 | 22.0 | 10.9 | 10.9 | 1713 | 3484 | 0.84 |
| 2 | 10.0 | 20.0 | 20.0 | 73.8 | 86.1 | -2.9 | 14.1 | 11.6 | 15.9 | 1722 | 3501 | 0.79 |
| 3 | 5.9 | 11.9 | 8.4 | 65.6 | 80.8 | 16.7 | 13.9 | 9.3 | 12.5 | 1480 | 3016 | 0.66 |
| 4 | 8.3 | 16.5 | 11.7 | 77.9 | 78.5 | 13.2 | 12.3 | 9.5 | 8.2 | 1332 | 2721 | 0.82 |
| 5 | 4.7 | 8.5 | 9.3 | 65.5 | 85.2 | 17.8 | 10.9 | 8.6 | 12.0 | 1428 | 2913 | 0.61 |
| 6 | 5.1 | 10.3 | 10.3 | 114.3 | 84.2 | 4.9 | 19.0 | 7.3 | 11.2 | 1415 | 2887 | 0.72 |
| 7 | 4.9 | 9.8 | 9.8 | 75.4 | 76.9 | 9.1 | 10.7 | 9.8 | 8.1 | 1386 | 2829 | 0.79 |
| 8 | 10.0 | 20.0 | 10.0 | 55.5 | 72.4 | -27.1 | 19.9 | 17.8 | 19.6 | 1742 | 3542 | 0.49 |
| 9 | 10.0 | 20.0 | 10.0 | 51.9 | 82.1 | -7.6 | 14.6 | 12.1 | 11.5 | 1563 | 3183 | 0.74 |
| 10 | 6.5 | 13.0 | 12.6 | 70.9 | 78.9 | 8.3 | 14.4 | 9.8 | 13.6 | 1781 | 3621 | 0.68 |
| 11 | 6.9 | 13.9 | 13.9 | 90.1 | 85.1 | 7.8 | 12.7 | 9.8 | 11.5 | 1488 | 3034 | 0.80 |
| 12 | 8.3 | 14.2 | 9.9 | 79.9 | 82.4 | -13.1 | 12.7 | 11.5 | 16.7 | 1410 | 2877 | 0.55 |
| 13 | 8.8 | 17.6 | 17.6 | 59.4 | 78.5 | 62.4 | 30.0 | 12.8 | 23.3 | 1741 | 3539 | 0.44 |
| 14 | 9.3 | 18.6 | 16.3 | 64.9 | 84.1 | 0.1 | 13.2 | 9.4 | 14.3 | 1521 | 3098 | 0.75 |
| 15 | 9.6 | 19.3 | 11.7 | 55.8 | 82.3 | 11.5 | 16.1 | 12.0 | 10.3 | 1550 | 3157 | 0.75 |
| 16 | 9.5 | 19.0 | 16.1 | 80.7 | 80.3 | 15.4 | 16.5 | 12.0 | 12.6 | 1761 | 3580 | 0.71 |
| 17 | 5.0 | 10.1 | 9.3 | 28.5 | 75.9 | 45.7 | 12.9 | 10.5 | 12.6 | 1487 | 3032 | 0.54 |
| 18 | 5.7 | 11.4 | 11.4 | 97.6 | 70.7 | 36.5 | 14.7 | 12.4 | 13.6 | 1385 | 2826 | 0.50 |
| 19 | 10.0 | 20.0 | 17.8 | 64.1 | 81.9 | -0.8 | 20.1 | 11.9 | 13.7 | 1625 | 3307 | 0.73 |
| 20 | 9.9 | 19.8 | 9.9 | 45.6 | 83.3 | -4.6 | 19.2 | 15.3 | 11.7 | 1646 | 3349 | 0.65 |
| mean | 7.9 | 15.7 | 12.7 | 68.1 | 80.8 | 10.3 | 16.0 | 11.2 | 13.2 | 1559 | 3175 | mdn 0.72 |
| SD | 2.0 | 4.1 | 3.5 | 19.1 | 4.2 | 19.6 | 4.5 | 2.3 | 3.5 | 142 | 286 | IQR 0.16 |

Figure 4. illustrates how well the predictions of the model matched each representative participant’s behavior on five unique target postures. For visualization purposes, we present the model components as stick figures in Cartesian coordinates. Each plot includes the likelihood (i.e., the target posture; black line), the arm posture prior (red line), and the response distribution (blue line). The actual the measured responses can be seen as purple dashed lines. As can be seen, the joints are perceived as more flexed when the prior is more flexed than the target posture, and more extended when the prior is more extended than the target posture. Responses are less deviant when the target posture is close to or overlaps with the prior.

**Figure 4.**
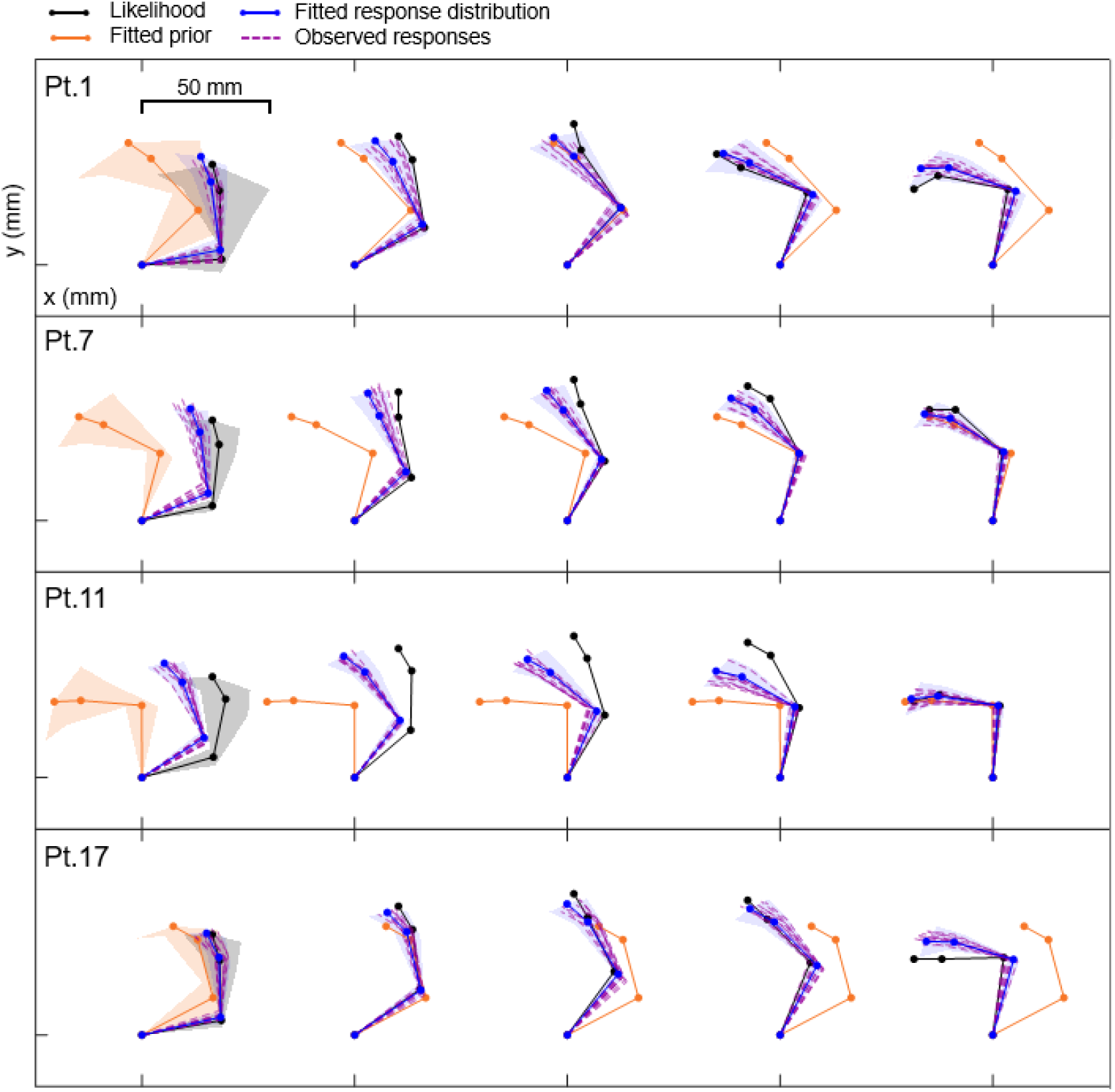
Model predictions in four participants plotted as stick figures in Cartesian coordinates. Modelled response (related to model’s posterior, in blue), the measured responses (in purple, dashed), target posture (i.e., the likelihood, in black) and the best-fitting prior of arm posture (in red) plotted for five unique target postures. For clarity, variability of prior and likelihood only shown for the most leftward configuration.

**Figure 5A** presents the best-fit prior means in Cartesian coordinates of all participants, which are relatively consistent across participants. Their mean (in green) yield the following angles (mean± SD, range): shoulder (68.1°±19.1°, [45.3°, 114.3°]), elbow (80.8°±4.2°, [70.7°, 86.1°]) and wrist (10.3°±19.7°, [-27.1°, 62.4°]).

**Figure 5.**
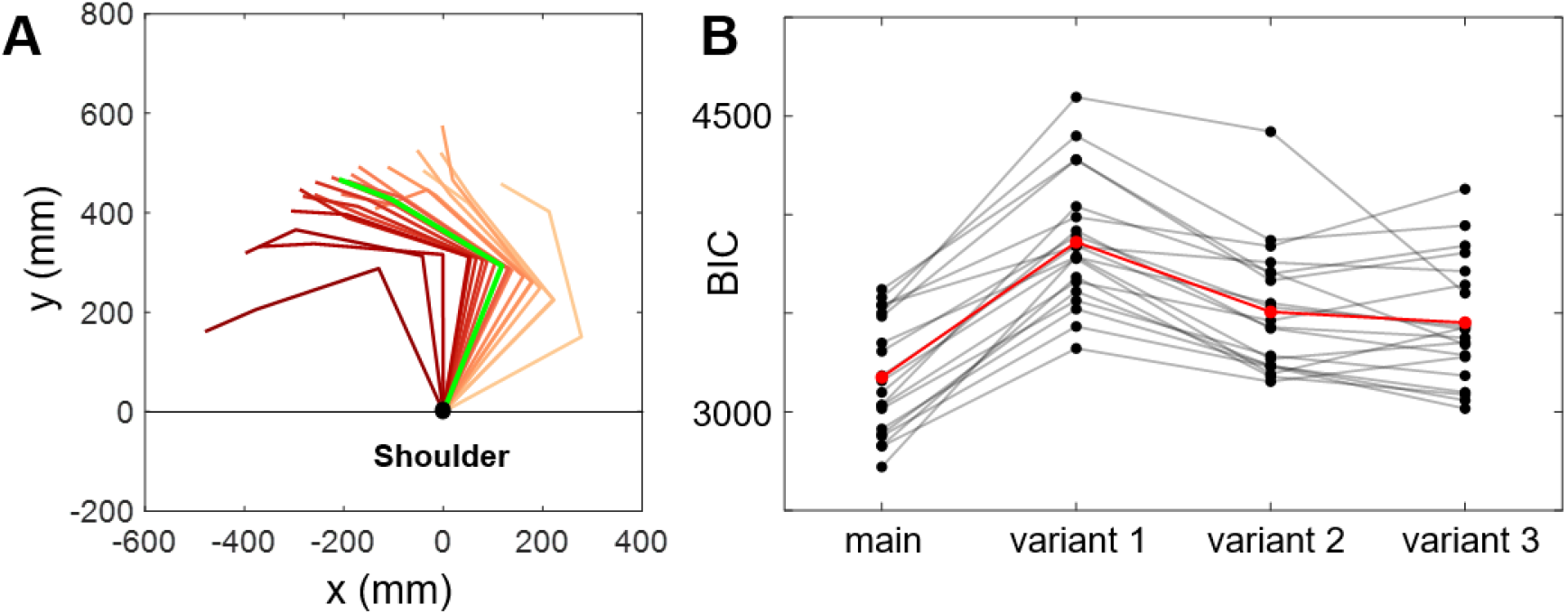
Best-fit priors and model comparison. **A**. Fitted priors plotted in Cartesian coordinates for all participants. The green line represents average fitted prior. **B**. BIC values for each participant and tested model (main: Bayesian model including non-uniform prior; variant 1: model variation without prior; variant 2: model variation with mean target posture as a prior; variant 3: model variation with starting position as a prior). Red line represents average BICs.

We next compared our best-fit model to three alternative variants: variant 1: uniform prior; variant 2: mean target posture prior; variant 3: starting posture prior (see Methods). The main model outperformed all three model variants (**Figure 5B**), as demonstrated by significantly lower BICs. The delta BIC between each model variant and the main model was greater than 30 for all participants.

## Discussion

The brain’s ability to estimate the position and orientation of the limbs from somatic signals is crucial for sensorimotor processing and behavior. Here, we formalized a computational model of arm proprioception based on Bayesian principles, which have been successfully applied before in somatosensation (25, 29). The computational process we propose estimates the perceived arm joint angle (shoulder, elbow, and wrist) by combining sensory likelihood with a postural prior. The key prediction of the model is that biases in arm posture perception, which vary as a function of arm posture, are due to the presence of a postural prior.

Our results strongly support this model. In all participants, the measured biases in arm posture perception varied as a function of the tested postures. Importantly, biases were consistently directed towards specific values in joint space, suggesting the involvement of a postural prior. Consistent with previous observations (6, 9) and the predictions of our Bayesian model, participants perceived their arm as too flexed when their physical arm was more extended than the prior posture (i.e., positive bias) and too extended when their physical arm was more flexed than the prior posture (i.e., negative bias). The bias decreased for postures that overlapped or were close to the inferred prior posture. The fitting results clearly corroborate this observation, showing that the main model outperformed a model variant without such best-fit prior (i.e. variant 1: uniform prior model). If the prior for arm posture were uniform, we would expect that matching responses would be unbiased; this was clearly not observed in our experiment.

Model fitting yielded an average prior with orientation of about 68°, 81°, and 10° for the shoulder, elbow, and wrist, respectively. These orientations correspond to a semi-flexed posture (see **Figure 5A**), with angles near to the middle of the anatomical range of joint motion (28). This prior also explains why previous studies show smaller proprioceptive matching errors for semi-flexed postures (6, 8), or perceptual biases towards such postures (9–11, 17). The influence of a semi-flexed prior posture on arm posture perception also fits very well with previous research reporting a systematic tendency to estimate the hand as closer to the torso and to the midline (e.g., (30–32).

Furthermore, our results are in line with previous research that used local anesthesia to study posture perception in absence of sensory signals. From a Bayesian standpoint, postural estimates in this case would mostly rely on the stored prior, due to the suppression of sensory input. In support, participants showed a strong and consistent estimation bias towards semi-flexed postures: when the elbow and wrist were extended, they were estimated more flexed, and vice versa (33–35).

While modeled priors are distributed quite homogeneously around a semi-flexed posture, we still observed some inter-individual differences. For instance, for the shoulder alone, the standard deviation for the prior mean across participants was ∼20°, with priors ranging from ∼28° to ∼114°. Such interindividual variation in prior posture likely explains the idiosynchraticity reported in endpoint (i.e., hand) localization estimates (36–39).

An important question is whether the best-fit priors have functional meaning. From a Bayesian perspective, the prior may represent an a-priori guess of arm posture, perhaps derived from previously adopted orientations, which would be combined with proprioceptive signals to increase perceptual precision. We also tested two alternative explanations. In one variant of the model (model variant 2) we assumed the average orientation of the target postures as the prior in the Bayesian computation. In another variant (model variant 3) we set the starting position, which is repeated at each trial, as prior. We found that the main model outperformed both these model variants. This demonstrates that matching errors were not merely directed to the average target posture, nor to the starting posture. This suggests that the prior we estimated through model fitting is not built up over the short time-range of the task, nor modulated based on the task’s contextual features—it is more likely a rather stable representation, perhaps reflecting the longer-term statistics of natural arm postures.

The suggested effect of a prior built upon statistical regularities in posture is in line with previous work linking natural statistics of arm and hand movements to motor biases (18, 40). For instance, previous research tracked mouse movements of computer-users for several days during their working hours, showing that movement direction spontaneously clustered at the cardinal axes. Such non-uniform directional distribution was modeled as prior, which well predicted directional biases in movement initiation, i.e., participants tend to initiate movements in the directions that are most represented in their own natural statistics (40). Future research is necessary to examine the origin of the postural prior; for instance by studying whether fitted priors at the individual level reflect statistical regularities in arm postures measured using wearable motion-tracking devices (e.g., 18, 41).

Another interesting finding pertains to the likelihood and prior variability estimated via model fitting. The likelihood variability was consistently smaller for the shoulder compared to the other joints in all participants, whereas the opposite pattern was observed for the prior, with wider priors for the shoulder joint, especially compared to the elbow joint (see **Table 2**). This may suggest that postural estimates for the shoulder relied more on sensory signals than on a-priori information. It follows that systematic errors were overall smaller for the shoulder compared to the elbow, which would be consistent with previous observations (e.g., 8, 42). Higher accuracy in shoulder position estimation may bring a functional advantage, because of noise propagation in a kinematic chain, the smaller the estimation errors at the level of the shoulder the more precise the control of hand position.

Here we specified a model that operates in angle-based coordinates, with outcomes that correspond to the requirements of the behavioral task, i.e., angular alignment of joints with visual stimuli, and fitted measurements, i.e., angular responses. To describe outcomes of other proprioception tasks, such as hand position localization, the model would need to account for the fact that errors propagate from their respective sources along the kinematic chain, from the shoulder to the hand. That is, while the estimated location of the elbow only depends on the measured shoulder angle and its variability, the location estimate of the hand depends on a combination of shoulder, elbow and wrist angles and their variability (e.g., 20, 25). For that, the model needs to be extended to include the reference frame transformations (geometric transformation of signals from angular to space-based coordinates).

From another perceptive, previous research has argued that different stages of reach planning rely on signals encoded in different coordinate systems: planning the initial movement direction is based on signals encoded in space-based coordinates such as vision, whereas converting the movement vector into motor commands relies on information encoded in angle-based coordinates such as proprioception (43, 44). An important question for future research is how postural priors come into play at different stages of motor planning. Does the angle-based prior we describe here bring information mostly relevant to define motor commands changing muscular torque? Can it also be flexibly transformed into space-based coordinates (e.g., in 25) contributing to the definition of the initial motor vector. These are interesting future directions that arise from this work.

## Acknowledgements

We extend our gratitude to the SensorimotorLab members for the numerous scientific discussions, and the Technical Support Group of DCC for their invaluable assistance and expertise.

V.C.P is supported by the Radboud Excellence Initiative grant. L.E.M is supported by the following grants: ERC 101076991 SOMATOGPS and NWO-VI.VIDI.221G.02. W.P.M. is supported by the following grants: NWA-ORC-1292.19.298, NWO-SGW-406.21.GO.009 and Interreg NWE-RE:HOME.

## Author contribution

All authors conceived and designed the study and interpreted the results; M.G.A.J. performed the experiment; V.C.P. and M.G.A.J. analyzed the data and prepared figures; V.C.P. drafted the manuscript; L.E.M and W.P.M. edited and revised the manuscript; W.P.M. supervised the project and provided resources.

## Data availability statement

Requests for data and code should be directed to and will be fulfilled by the lead contact, Valeria Peviani.

## Supplemental material

**Supplemental Figure 1.**
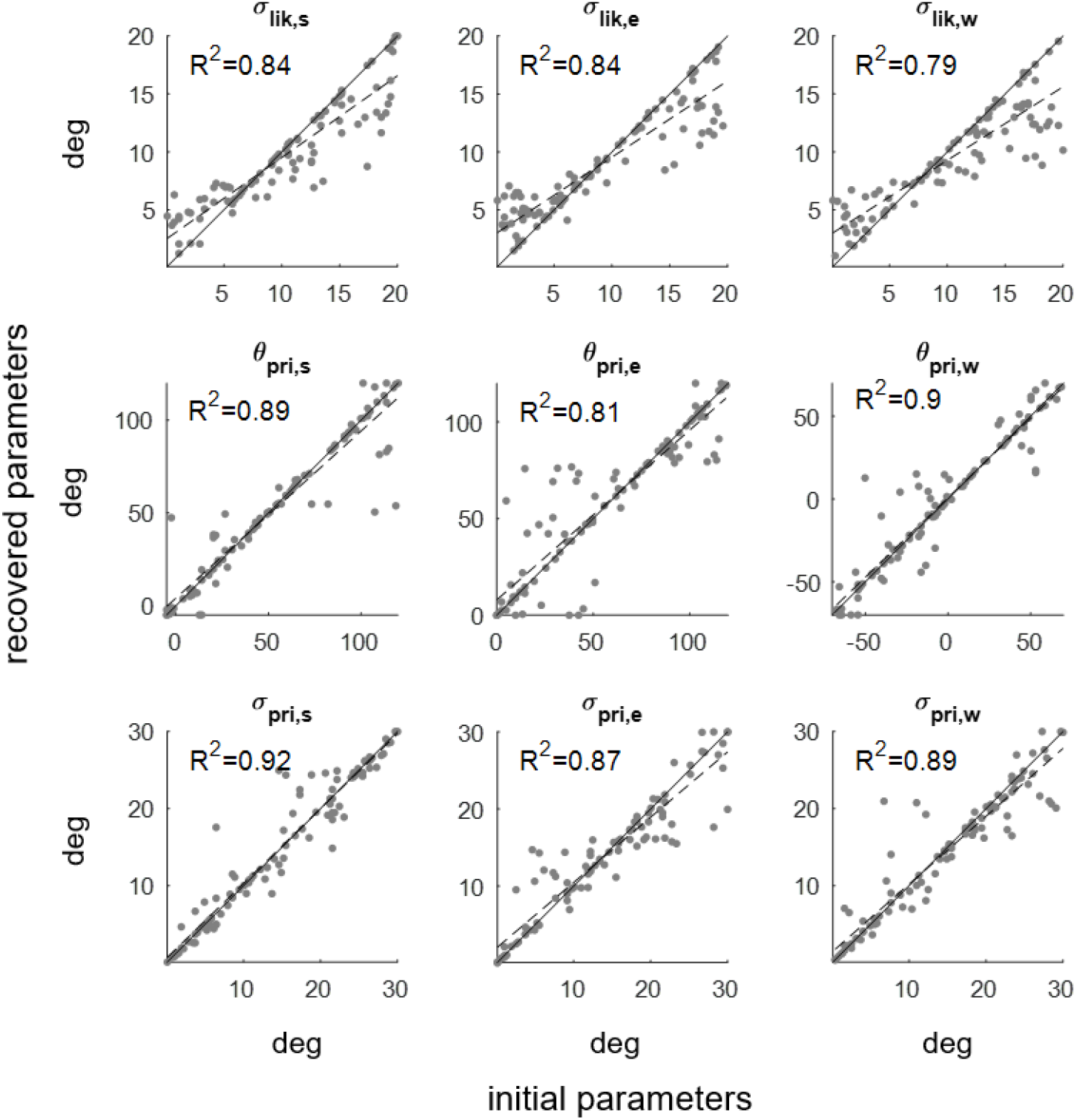
Parameters recovery analysis. Predicted parameter values plotted against the initial parameter values for the nine free parameters of the main model. The grey solid line represents the diagonal, whereas the dashed line shows the regression line.

## References

1. Héroux ME, Butler AA, Robertson LS, Fisher G, Gandevia SC. Proprioception: a new look at an old concept. J Appl Physiol 132 American Physiological Society Rockville, MD: 811–814, 2022.

2. Proske U, Gandevia SC. The Proprioceptive Senses: Their Roles in Signaling Body Shape, Body Position and Movement, and Muscle Force. Physiol Rev 92: 1651–1697, 2012. doi: 10.1152/physrev.00048.2011.-This.

3. Shadmehr R, Mussa-Ivaldi S. Biological learning and control: how the brain builds representations, predicts events, and makes decisions. Mit Press, 2012.

4. Oostwoud Wijdenes L, Medendorp WP. State estimation for early feedback responses in reaching: intramodal or multimodal? Front Integr Neurosci 11: 38, 2017.

5. Proske U, Chen B. Two senses of human limb position: methods of measurement and roles in proprioception. Exp Brain Res 239: 3157–3174, 2021.

6. Fuentes CT, Bastian AJ. Where is your arm? Variations in proprioception across space and tasks. J Neurophysiol 103: 164–171, 2010.

7. Klein J, Whitsell B, Artemiadis PK, Buneo CA. Perception of arm position in three-dimensional space. Front Hum Neurosci 12: 331, 2018.

8. King J, Harding E, Karduna A. The shoulder and elbow joints and right and left sides demonstrate similar joint position sense. J Mot Behav 45: 479–486, 2013.

9. Abi Chebel NM, Roussillon NA, Bourdin C, Chavet P, Sarlegna FR. Joint specificity and lateralization of upper limb proprioceptive perception. Percept Mot Skills 129: 431–453, 2022.

10. Chen B, Allen T, Proske U. Position sense at the human forearm over a range of elbow angles. Exp Brain Res 239: 675–686, 2021.

11. Goble DJ, Brown SH. Upper limb asymmetries in the perception of proprioceptively determined dynamic position sense. J Exp Psychol Hum Percept Perform 36: 768, 2010.

12. Bergenheim M, Ribot-Ciscar E, Roll J-P. Proprioceptive population coding of two-dimensional limb movements in humans: I. Muscle spindle feedback during spatially oriented movements. Exp Brain Res 134: 301–310, 2000.

13. Rossetti Y, Meckler C, Prablanc C. Is there an optimal arm posture? Deterioration of finger localization precision and comfort sensation in extreme arm-joint postures. Exp Brain Res 99: 131–136, 1994.

14. Yamamoto S, Kitazawa S. Reversal of subjective temporal order due to arm crossing. Nat Neurosci 4: 759–765, 2001.

15. Tamè L, Azañón E, Longo MR. A conceptual model of tactile processing across body features of size, shape, side, and spatial location. Front Psychol 10: 423359, 2019.

16. Badde S, Heed T. The hands’ default location guides tactile spatial selectivity. Proceedings of the National Academy of Sciences 120: e2209680120, 2023.

17. Gritsenko V, Krouchev NI, Kalaska JF. Afferent input, efference copy, signal noise, and biases in perception of joint angle during active versus passive elbow movements. J Neurophysiol 98: 1140–1154, 2007.

18. Howard IS, Ingram JN, Körding KP, Wolpert DM. Statistics of natural movements are reflected in motor errors. J Neurophysiol 102: 1902–1910, 2009.

19. Faisal AA, Selen LPJ, Wolpert DM. Noise in the nervous system. Nat Rev Neurosci 9: 292–303, 2008.

20. Oh K, Prilutsky BI. Transformation from arm joint coordinates to hand external coordinates explains nonuniform precision of hand position sense in horizontal workspace. Hum Mov Sci 86: 103020, 2022.

21. Van Beers RJ, Sittig AC, Denier van der Gon JJ. The precision of proprioceptive position sense. Exp Brain Res 122: 367–377, 1998.

22. Ma WJ, Kording KP, Goldreich D. Bayesian models of perception and action: An introduction. MIT press, 2023.

23. Pouget A, Beck JM, Ma WJ, Latham PE. Probabilistic brains: knowns and unknowns. Nat Neurosci 16: 1170–1178, 2013.

24. Clemens IAH, de Vrijer M, Selen LPJ, van Gisbergen JAM, Medendorp WP. Multisensory processing in spatial orientation: An inverse probabilistic approach. Journal of Neuroscience 31: 5365–5377, 2011. doi: 10.1523/JNEUROSCI.6472-10.2011.

25. Peviani VC, Miller LE, Medendorp WP. Biases in hand perception are driven by somatosensory computations, not a distorted hand model. Current Biology 34: 2238-2246.e5, 2024. doi: 10.1016/j.cub.2024.04.010.

26. Oldfield RC. The assessment and analysis of handedness: the Edinburgh inventory. Neuropsychologia 9: 97–113, 1971.

27. Veale JF. Edinburgh Handedness Inventory - Short Form: A revised version based on confirmatory factor analysis. Laterality 19: 164–177, 2014. doi: 10.1080/1357650X.2013.783045.

28. Aizawa J, Masuda T, Hyodo K, Jinno T, Yagishita K, Nakamaru K, Koyama T, Morita S. Ranges of active joint motion for the shoulder, elbow, and wrist in healthy adults. Disabil Rehabil 35: 1342–1349, 2013.

29. Miller LE, Fabio C, Azaroual M, Muret D, van Beers RJ, Farn A, Pieter Medendorp W. A neural surveyor to map touch on the body. PNAS 119: 1–12, 2022. doi: 10.1073/pnas.2102233118/-/DCSupplemental.

30. Haggard P, Newman C, Blundell J, Andrew H. The perceived position of the hand in space. Percept Psychophys 62: 363–377, 2000.

31. Wilson ET, Wong J, Gribble PL. Mapping proprioception across a 2D horizontal workspace. PLoS One 5: e11851, 2010.

32. Peviani V, Bottini G. Proprioceptive errors in the localization of hand landmarks: What can be learnt about the hand metric representation? PLoS One 15: e0236416, 2020. doi: 10.1371/JOURNAL.PONE.0236416.

33. Melzack R, Bromage PR. Experimental phantom limbs. Exp Neurol 39: 261–269, 1973.

34. Inui N, Masumoto J, Ueda Y, Ide K. Systematic changes in the perceived posture of the wrist and elbow during formation of a phantom hand and arm. Exp Brain Res 218: 487–494, 2012.

35. Inui N, Walsh LD, Taylor JL, Gandevia S. Dynamic changes in the perceived posture of the hand during ischaemic anaesthesia of the arm. J Physiol 589: 5775–5784, 2011.

36. Rincon-Gonzalez L, Buneo CA, Helms Tillery SI. The proprioceptive map of the arm is systematic and stable, but idiosyncratic. PLoS One 6: e25214, 2011.

37. Kuling IA, Van Der Graaff MCW, Brenner E, Smeets JBJ. Matching locations is not just matching sensory representations. Exp Brain Res 235: 533–545, 2017.

38. Liu Y, Sexton BM, Block HJ. Spatial bias in estimating the position of visual and proprioceptive targets. J Neurophysiol 119: 1879–1888, 2018.

39. Wang T, Zhu Z, Kana I, Yu Y, He H, Wei K. Accuracy of hand localization is subject-specific and improved without performance feedback. Sci Rep 10: 19188, 2020.

40. Slijper H, Richter J, Over E, Smeets J, Frens M. Statistics Predict Kinematics of Hand Movements During Everyday Activity. J Mot Behav 41: 3–9, 2009. doi: 10.1080/00222895.2009.10125922.

41. Ingram JN, Körding KP, Howard IS, Wolpert DM. The statistics of natural hand movements. Exp Brain Res 188: 223–236, 2008. doi: 10.1007/s00221-008-1355-3.

42. Galofaro E, D’Antonio E, Patané F, Casadio M, Masia L. Three-Dimensional Assessment of Upper Limb Proprioception via a Wearable Exoskeleton. Applied Sciences 11, 2021. doi: 10.3390/app11062615.

43. Sober SJ, Sabes PN. Multisensory Integration during Motor Planning. The Journal of Neuroscience 23: 6982, 2003. doi: 10.1523/JNEUROSCI.23-18-06982.2003.

44. Sober SJ, Sabes PN. Flexible strategies for sensory integration during motor planning. Nat Neurosci 8: 490–497, 2005. doi: 10.1038/nn1427.

